# PhylUp: phylogenetic alignment building with custom taxon sampling

**DOI:** 10.1101/2020.12.21.394551

**Authors:** Martha Kandziora

## Abstract

In recent years it has become easier to reconstruct large-scale phylogenies with more or less automated workflows. However, they do not permit to adapt the taxon sampling strategy for the clade of interest. While most tools permit a single representative per taxon, PhylUp – the workflow presented here - enables to use different sampling strategies for different taxonomic ranks, as often needed for molecular dating analyses or for a large outgroup sampling. While PhylUp focuses on user-defined sampling strategies, it also facilitates the updating of alignments with new sequences from local and online sequence databases and their incorporation into existing alignments. To start a PhylUp run at least one sequence per locus has to be provided, PhylUp then adds new sequences to the existing one by internally using BLAST to find similar sequences and filters them according to user settings. Taxonomic sampling is increased compared to available tools and the custom taxonomic sampling allows to use automated workflows for new research fields. The workflow is presented in detail and I demonstrate the usability.

## 1 Introduction

Knowledge of the relationships of species is important for various fields of research (Felsenstein, 1985). Given the large number of new sequences that are published daily, for example in the GenBank database (Benson et al., 2018) by the National Center for Biotechnology Information (NCBI), any published organismal phylogeny is quickly outdated. Regularly checking for new sequences for taxa of interest and updating the corresponding alignments and phylogenies is time-consuming, if done at all. In recent years several new methods have been developed that can be used to infer alignments and phylogenies using automated workflows (Antonelli et al., 2017; Bennett et al., 2018; Drori et al., 2018; Izquierdo-Carrasco et al., 2014; Portik and Wiens, 2020; Shipunov, 2020; Smith and Walker, 2019). Those tools differ in aspects of the alignment building and phylogenetic reconstruction processes, but are limited to calculate phylogenies based on one sample per species or genus (depending on the tool).

For many research questions however, researchers require more flexible sampling strategies. Studies that are interested in confirming species monophyly need several samples per species. Broad-scale analyses often need larger outgroup sampling, while a single individual might be sufficient for the clade of interest. Datasets for molecular dating often consist of a sparse sampling with a few representatives for all subfamilies within a family that allows to apply fossil and secondary age calibrations, but with a better representation for the clade of interest.

Here, I present – PhylUp – a new python package to automatically build or update alignments based on complex, flexible-taxon sampling strategies. PhylUp is also able to include unpublished locally available sequences to the alignment.

## 2 Workflow

PhylUp is a command-line tool written in python3. Internally, the tool depends on other programs to retrieve additional sequences to build the alignment and to calculate a phylogenetic tree: 1) The Basic Local Alignment Search Tool v. 2.9.0 from ncbi (BLAST; Camacho et al., 2009) searches for similar sequences to the input sequences in a database; 2) PaPaRa v. 2.5 (Berger and Stamatakis, 2011), an alignment program which uses a reference alignment and reference phylogeny to add new sequences to the alignment or MAFFT v. 7.45 (Katoh and Standley, 2013); 3) RAxML-NG v. 0.9.0 (Kozlov et al., 2019) as phylogenetic tree inference program; 4) EPA-NG v. 0.3.5 (Barbera et al., 2019) to place new sequences into an existing phylogeny; and 5) Modeltest-NG v. 0.1.6 (Darriba et al., 2019) to infer the best fitting substitution model.

Figure 2.1 presents the general workflow of PhylUp. A single-locus alignment or a single sequence serves as input. Additionally, a translation table connecting input sequence names with NCBI taxon names, and a configuration file have to be provided (Fig. 2.1). Optional, the user can provide a phylogeny, or a list of GenBank accessions to be excluded. Sampling of alignment and phylogeny do not need to match, which ensures that phylogenies produced with a concatenated dataset can be used to update the single-gene alignments. The workflow removes empty sequences and additional tips in the tree prior to the update.

**Figure 2.1:**
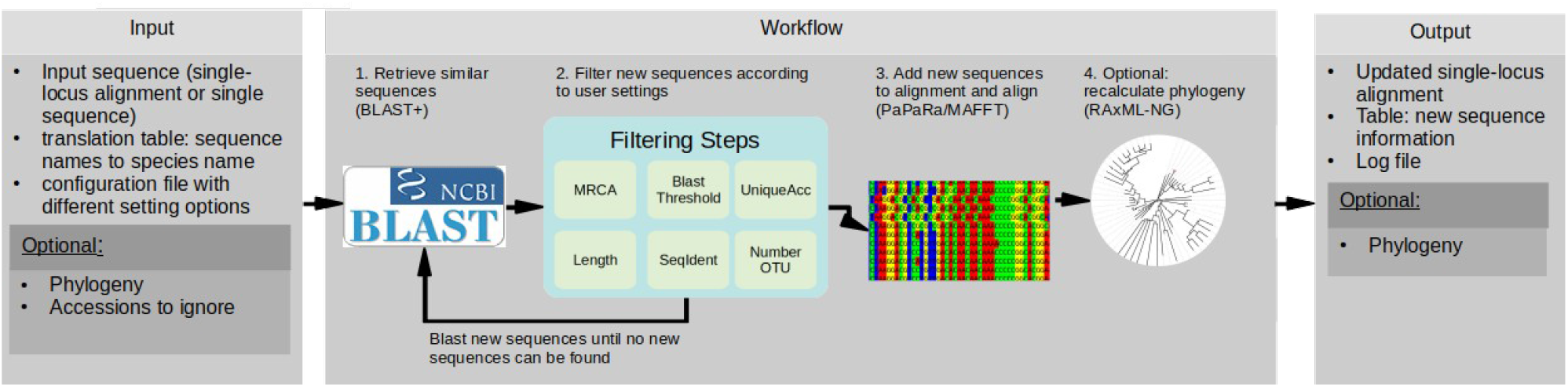
Overview of PhylUp workflow. Initial sequence(s) are blasted, filtered according to user settings and then assembled. Optionally, a Maximum Likelihood tree is calculated.

Step 1 of the workflow blasts the input sequence(s) against a local copy of the GenBank database (Benson et al., 2018), a local sequence database (with, for example, unpublished new sequences) or a combination of both. The BLAST results are only limited by the number of sequences returned by the search (hitlist_size). The BLAST step uses user-provided similarity values (e-value) to retrieve similar sequences from the database. Thus, the few numbers of incorrectly identified taxa, and ambiguous sequence from GenBank annotation should not be an issue (Leray et al., 2019).

Step 2 filters BLAST results according to user settings: a) only sequences below the defined BLAST e-value are retained, b) sequences that differ more than a user-defined length threshold compared to the input sequences are removed, c) sequences not part of the most recent common ancestor (mrca) are removed, d) optional: only sequences that differ from existing ones are kept – at least one point mutation for same taxon identifier (including synonyms), e) if multiple sequences per rank are available, new OTU will be preferred (optional), f) if more sequences were found per OTU as defined by the user a subset is selected. Subset selection is done either by sequence length or another local BLAST search. All new sequences passed the different filtering steps are blasted again until no new sequences can be found or until the maximal number of BLAST rounds is reached (status_end setting).

Step 3, single sequence provided as input: initial alignment is generated with MAFFT. An alignment is provided as input: Newly retrieved and filtered sequences are added to the alignment using PaPaRa.

Step 4 (optional) calculates a phylogeny. New sequences are placed into an existing phylogeny using EPA-NG using information based on the reference alignment and reference tree (Barbera et al., 2019). A user-provided phylogeny can serve as a reference tree, if not PhylUp uses a fully resolved tree generated by Modeltest-NG. Then, EPA-NG is used to place the new sequences into the phylogeny using information based on the reference alignment and tree. The algorithm odoes not update the internal relationships. The resulting phylogeny serves as a starting tree for the phylogenetic reconstruction using RAxML-NG.

Further settings are available via the configuration file (see Supplement 1; Wiki at https://github.com/mkandziora/PhylUp). All internal translation between taxon names, taxon identifiers and hierarchical information, which is based on the taxonomic information provided by NCBI are performed by the python package ncbiTAXONparser a flexible wrapper to retrieve ncbi taxonomic information (https://github.com/mkandziora/ncbiTAXONparser).

### 2.1 Options for custom taxon sampling strategies across different hierarchical levels

To allow for different taxonomic sampling strategies, PhylUp limits the number of sequences per operational taxonomic unit (OTU) and taxonomic rank based on user settings. If the initial BLAST step finds more sequences per taxon than defined by the threshold value, there are two options available on how to select the defined number of sequences per OTU. First, the program selects the longest sequences, or second, a second local BLAST search is conducted. If a second BLAST search is used for the filtering, a sequence already present in the alignment or a randomly chosen sequence from the newly found ones will be used to blast against all other sequences from the locus with the same NCBI taxon ID (either based on the ID of the sequence or on the rank filtering option - see below).

The local BLAST search reveals a proxy for sequence divergence across the same OTU. As a first step, any sequence that does not fall within the mean plus or minus standard deviation of sequence similarity will be filtered out (see Supplement 7.1.4). If several sequences are available, it is likely that mis-identifications are filtered out due to higher sequence dissimilarity compared to the remaining sequences. If only few sequences or multiple but similarly diverging sequences are available none will be filtered. Finally, from the sequences that pass the filtering criterion, sequences are randomly selected as representative.

Additionally to the species number filter, a rank filtering level can be defined. In this case, the filtering does not apply to the NCBI taxon ID, which can be of any taxonomic hierarchical level - usually the one provided by GenBank is used. Instead, by using the rank filter, the program can select a number of representatives of higher levels, e.g. by ‘genus’ or ‘tribe’. The rank delimitation follows the NCBI taxonomy. By using a combination of the species threshold and rank filters the user can build custom-sampled alignments. For example, as often done for molecular dating approaches, it is possible to update alignments to include two sequences per species for a genus, one sequence per genus for the respective tribe and then another two sequences for the different tribes of the family. During the filtering steps, all names that result in the same NCBI taxon ID are considered to be the same accepted taxon, thus synonyms are included under the same accepted taxon and if the filtering is done by genus, all taxa that are part of the genus are being considered. Further, if multiple representatives of a higher rank shall be selected, PhylUp will preferentially add OTU that have not yet been sampled.

When multiple loci shall be updated, PhylUp can match OTU across loci (using the run_multiple function).

### 2.2 Using different sequence databases

Instead of querying GenBank for new sequences, local sequences which have not yet been uploaded to GenBank can be added as well. Therefore, a folder with the sequences formatted in FASTA format and a table where sequence names correspond to species names is needed (see online repository for example files). PhylUp will generate a local BLAST database of those sequences and will add new sequences to the alignment if they pass the filtering criteria. Using a user-defined database can either be used as an additional step that happens before blasting against a local GenBank database or it can be used instead of a GenBank database (see Fig. 2.2).

**Figure 2.2:**
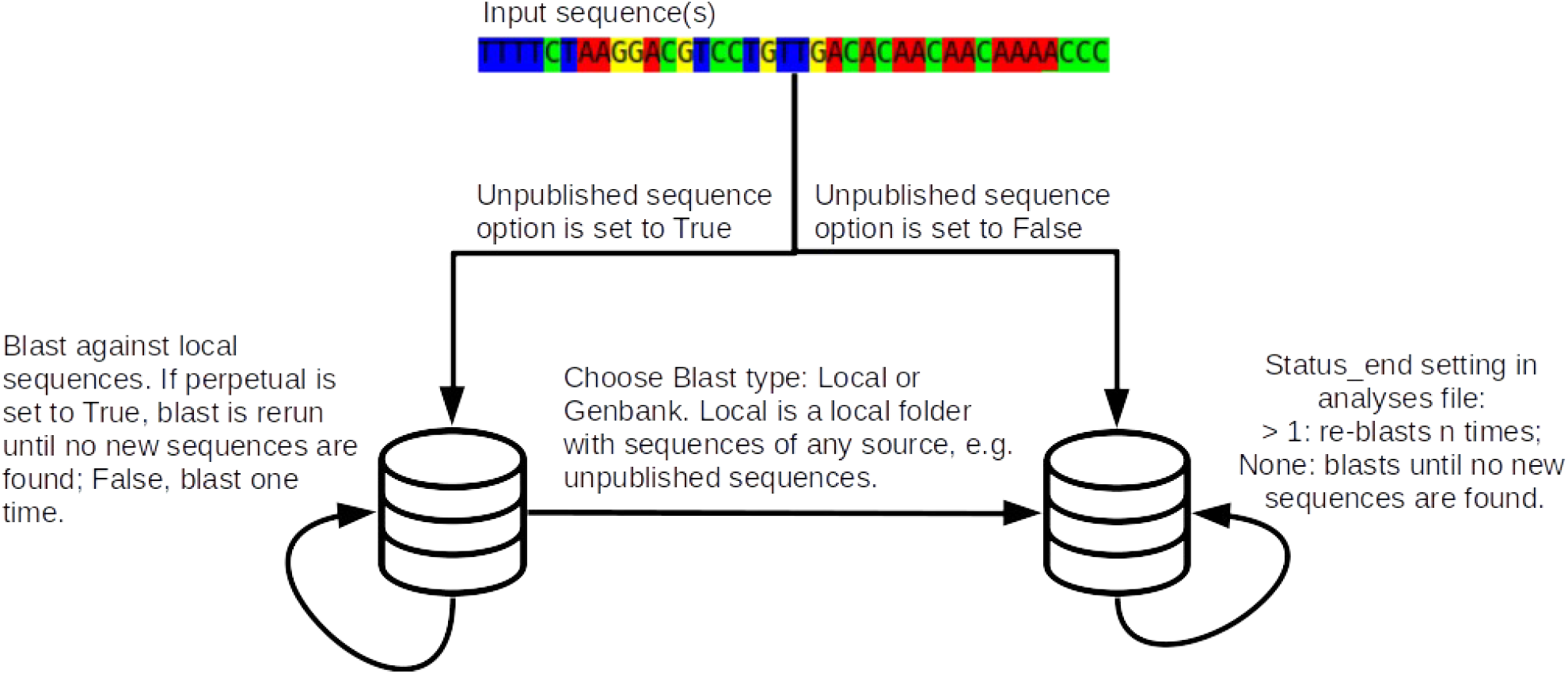
Overview of the different options for sequence databases and search strategies in PhylUp.

## 3 Generating custom taxon sampling alignments using PhylUp

To demonstrate PhylUp I used two examples: 1) one to show the usability during molecular dating: using different number of samples per OTU for three different rank filters across Rosaceae (for details see Supplement 1.2) to build a Rosaceae alignment with a focus on *Rosa*, but with samples across the subfamilies and 2) for comparison to other tools: a simple taxonomic update using Phylup (two samples per species) of a Senecioneae datasets (Kandziora et al., 2017, 2016). Taxon sampling was then compared to pyPHLAWD and OneTwoTree (see Supplementary Material 1.2 for more details (Drori et al., 2018; Smith and Walker, 2019)).

Both alignments generated with PhylUp are well sampled. Building a custom sampled alignment from scratch using PhylUp resulted in a phylogeny fulfilling the taxonomic sampling as set in the settings (Fig. 3.1). The Rosaceae alignment represents different taxonomic levels as needed for molecular dating analyses (Supplement 1.2). The updating of the Senecioneae alignments increased the amount of sequences and taxa across all used markers and adds more sequences and species as the two other tools (Fig. 3.2). The phylogenetic relationship of the updated phylogenies represent the current phylogenetic relationships (see Supplementary Material 1.3).

**Figure 3.1:**
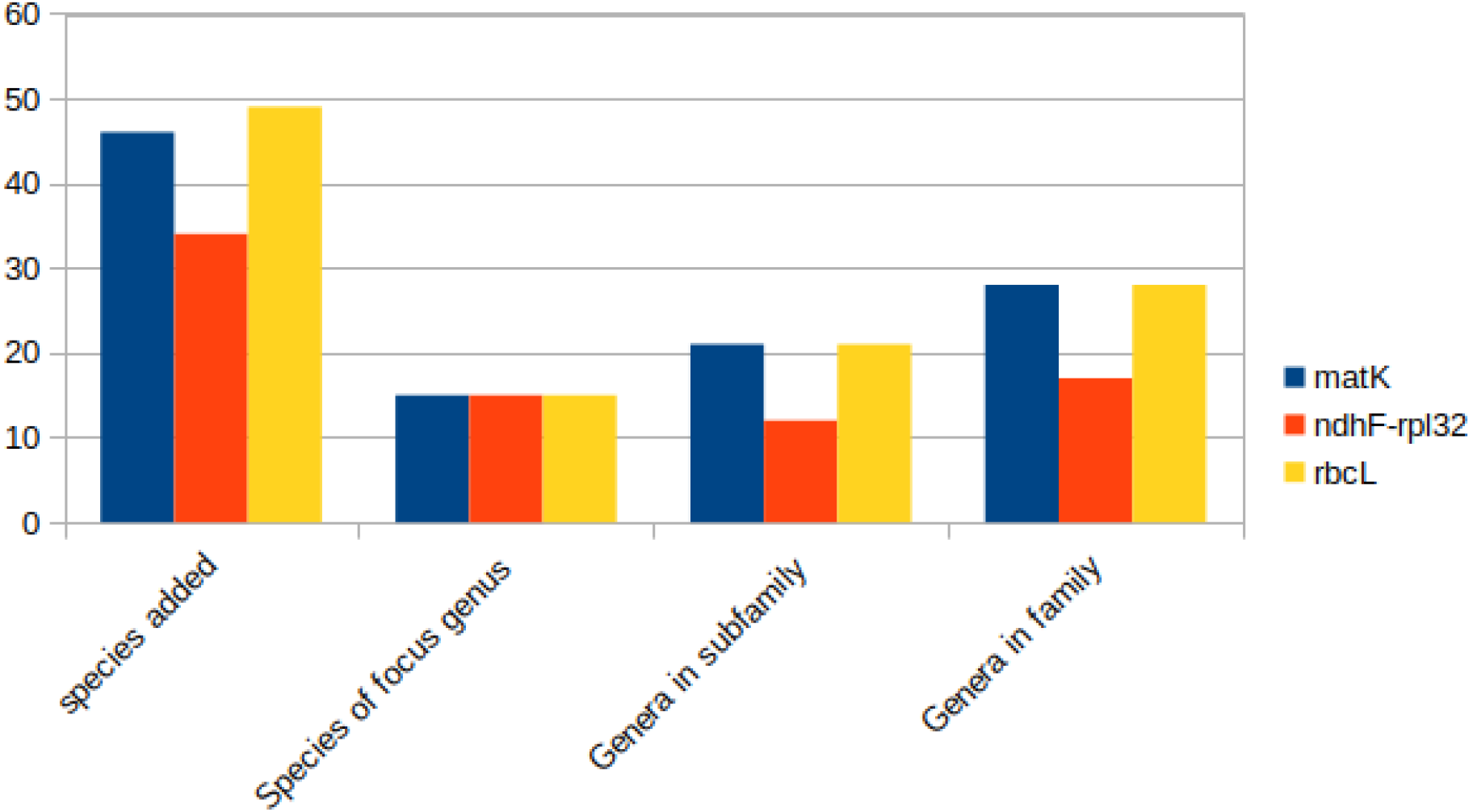
Sampling across different taxonomic ranks after using PhylUp to generate a Roaceae alignment.

**Figure 3.2:**
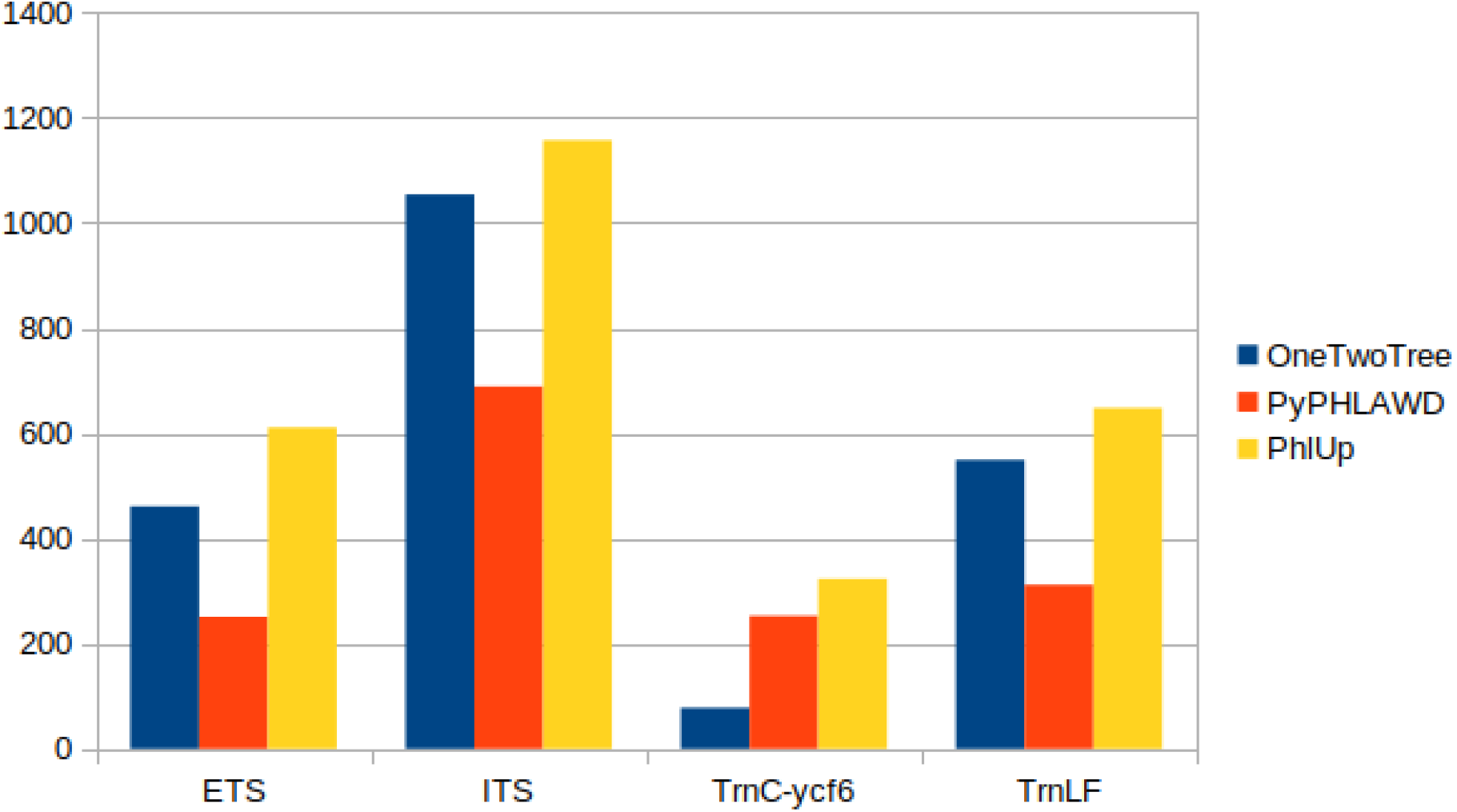
Species sampling across the different loci based on Senecioneae alignments in comparison to tools that build automatically phylogenies (alignments).

## 4 Discussion

PhylUp offers a user-friendly method to update alignments and customize the taxonomic sampling strategy. Several options set PhylUp apart from other tools to easily calculate phylogenies. The filtering of the available sequences to represent taxa by different amounts of sequences across different taxonomic hierarchies is a new feature among the different tools available today. Overall, the updated alignments using PhylUp are well sampled and minimize missing data while adding most taxa available on GenBank (Supplement 1.3).

Out of the suite of existing automated workflows, two new tools offer the possibility of adding all or multiple available sequences per taxon (Portik and Wiens, 2020; Sanchez Reyes et al., 2020). Nevertheless, none of them permit to define a certain number of species for higher taxonomic ranks. Furthermore, often researchers need to add new (unpublished) sequences into their alignments before calculating trees, which is only possible using one workflow currently (Portik and Wiens, 2020). Physcraper, a workflow developed by colleagues and me (Sanchez Reyes et al., 2020) can also update user-provided alignments utilizing BLAST, but each tool has a different focus: PhylUp concentrates on filtering sequences based on different taxonomic ranks while physcraper focuses on the integration with Open Tree of Life.

Despite existing automated tools to build alignments and phylogenies, many researchers still invest considerable extra time in manual alignment building and curation of matrices. There can be benefits to such an approach: Depending on the genetic marker, the alignment may be challenging due to repetitive sequences or indels, and hand-curated sequence alignments may provide optimal resolution for a clade of interest. PhylUp can take either a single species or an existing alignment similar to others as input and guidance for the alignment process of the new sequences (Izquierdo-Carrasco et al., 2014; Sanchez Reyes et al., 2020).

PhylUp is meant as an addition to already available tools that generate phylogenies from scratch. It is the first program that concentrates on a flexible custom taxonomic sampling strategy during alignment building. While permitting custom taxonomic sampling strategies, it additionally uniquely combines different options: adding sequences from unpublished sources, updating alignments and removing unwanted sequences. The tool is currently available via GitHub (https://github.com/mkandziora/PhylUp), including documentation and examples.

## 6 Acknowledgment

This workflow is based on ideas I developed during and after working in Emily Jane McTavish’s group on updating phylogenies using the Open Tree of Life framework. I would like to thank her for this opportunity. Thanks to the future reviewers for the time and suggestions for improvement.

## 7 Data availability

Results and data are available via Figshare at the point of publication. Example analysis files can also be found in the GitHub repository. All the other data is available via NCBI.

