## Supplementary Material for "PhylUp: phylogenetic alignment building with custom taxon sampling"

### 1 Supplement

#### 1.1 Detailed explanation of PhylUp options

##### 1.1.1 Defining the mrca

If the user wants to update the alignment either to beyond the current sampling of the data or only to a subclade of the input, they can define the mrca of the clade of interest in the configuration file by providing the NCBI taxon identifier. If the user provides no mrca, the tool calculates the mrca of all input sequences. Overall, the setting of the mrca needs to be handled with care. If the group of interest is not monophyletic but defined as being monophyletic in the NCBI taxonomy, closely related sequences are not added. It is possible to add several mrca (e.g. mrca = NCBI taxonid1, NCBI taxonid2, NCBI taxonid3) or to choose the next higher monophyletic node to account for this issue that is related to the ncbi taxonomic database. For example, the genus *Senecio* in its current circumscription contains genera that were not part of the genus before, plus many species that were defined to belong to the genus are actually only distantly related to the genus (Pelser et al., 2007). To focus on the genus *Senecio* in its current circumscription (*Senecio* s.str.), one can define all NCBI taxon IDs for the genera that are nested within *Senecio*: Hasteola Raf., *Lasiocephalus* Willd. ex Schltdl., *Iocenes* B.Nord., *Aetheolaena* Cass., *Culcitium* Humb. & Bonpl. and *Robinsonia* DC. This ensures to get all the sequences for the group of interest, but will also add *Senecio* species, that are actually only distantly related to *Senecio* s.str. and which need to be transferred to other genera or new yet undescribed genera.

Furthermore, the PhylUp settings allow the user to extend the input alignment to any clade of interest, which requests some attention to the resulting alignments by the user to ensure that the marker used is appropriate. If the sequences highly diverge from each other, the resulting alignment can be too variable to infer a robust phylogeny.

##### 1.1.2 Filtering to the correct number of sequences using BLAST

The sequences are being filtered according to bit-scores returned by BLAST. Bit-scores are a measure for sequence similarity. Bit-score is a log-scaled value of the score, which is a numerical value that describes the overall quality of an alignment. While scores are depending on database size, the re-scaled bit-scores do not (https://www.ncbi.nlm.nih.gov/BLAST/tutorial/Altschul-1.html). Sequences are being removed during the filtering of sequences if their bit-score is not within the mean plus/minus the standard deviation of sequence similarity in relation to the queried sequence.

Outlier sequences are likely mis-identification or mislabeling in GenBank. They are likely more diverging to the remaining sequences that they will be outside the allowed range of the mean and SD of sequence similarity. If within a taxon sequence divergence is large, sequences will be within the range as the mean and SD will be larger and allows to randomly pick sequences that represent the divergence.

##### 1.1.3 Further configuration settings

The program will delete sequences that are much shorter or longer than the alignment (as defined in min_len and max_len) and will prune the beginning and end of the alignment to have at least a certain amount of sequence information present (defined in trim_perc). These options are important if, for example, a database of plastome sequences was provided as input database, but the input sequences are shorter loci. Then, the max_len option allows to consider those long sequences, while trim_perc will ensure that the alignments are trimmed to represent the shorter loci. A further setting option is to use status_end in the analysis file to restrict the number of 'blast rounds'. This allows to limit the number of newly found sequences to the most similar once of the input.

##### 1.1.4 Concatenation of single-locus PhylUp runs

If single-locus alignments were updated, there is another python package available, Concat (https://github.com/mkandziora/phylogenetic_concatenation), that can combine the data and calculate a phylogeny from the concatenated data (see example files in the PhylUp package). The Concat package can concatenate sequences from the same OTU. Species that are only represented by a single marker are added as long as the amount of missing species is above a user-defined threshold (taxon missingness value). There is also an option available where the user can specify which sequences shall be concatenated by supplying a table with the corresponding accession numbers of sequences. This will allow the user to choose which of potentially multiple sequences per loci and taxa shall be concatenated – an advantage for example if the user knows that certain sequences are derived from the same DNA material. This option will grant even more flexibility but still minimizes steps that can easily be handled by the computer. In the end, a concatenated phylogeny can be calculated. Concat can be used independently from PhylUp, then the different alignments and a translation table of the sequence names to NCBI species names has to be provided.

#### 1.2 Build custom sampled alignments

##### 1.2.1 Datasets and Settings

For comparison to existing tools, I conducted a PhylUp run with simple taxonomic settings. I updated the *Senecio* and Senecioneae single-locus alignments from Kandziora et al. (2016) and Kandziora et al. (2017). I used the single-locus alignments of the nuclear ribosomal markers internal transcribed spacer 1 and 2 (ITS) and the external transcribed spacer (ETS), as well the plastid markers of *trn*L-*trn*F and *trn*C-*ycf*6. The mrca was defined as Senecioneae (NCBI taxon ID 102812). For all examples using PhylUp, I used a local GenBank database that was downloaded on October 17th, 2019.

Settings for the configuration are the following: For BLAST, the e-value is set to 0.001 and the hitlist size to 500. The settings for the alignment minimum length is set to 0.65%, maximum length to 2.5% and the ends are trimmed if less than 45% have no sequences available. The filtering of taxa is done using local blast and is limited to 1 sequences per species, i.e. subspecies and varieties are not all sampled with two sequences per OTU where available. I concatenated the chloroplast markers as well as the nuclear ribosomal markers separately. Plastid and nuclear markers were not concatenated due to known phylogenetic inconsistencies (Pelser et al., 2010). For the concatenation, I allowed a maximum of 25% missing taxa in the alignment.

To show how to build an alignment with custom taxonomic sampling settings, I initiated PhylUp to build a Rosaceae phylogeny. As input, I provided three single input sequences of *Rosa laevigata*: trnLF, ndhf-rpl32 intergenic spacer and rbcL. I used similar settings as above. I set the blast hitlist size to 1500 and set different settings for the number of sequences, the rank and the mrca: Sampling of taxa was set to sample a maximum of 15 sequence per genus for Rosa, two sequences per genus for the subfamily Rosoideae and two sequences per tribe for the family Rosaceae (see Table S1). Furthermore, I enabled the preferred taxon option to maximize sampling across loci. In order to update the alignment with different sampling strategies for different rank levels and across loci I used the run_multiple function available from PhylUp. To allow for the different filtering options, the different_level option has to be enabled in the configuration file of step two and three (see examples in the GitHub repository for more details). A concatenated phylogeny was calculated using RAXML-NG.

Table S1: Settings used to update alignments using different sampling strategies for different taxonomic ranks.

| Update level | Mrca (ncbi ID) | number per OTU | Rank level |
|---|---|---|---|
| 1 | Rosa (3764) | 15 | Genus |
| 2 | Rosoideae (171638) | 2 | Genus |
| 3 | Rosaceae (3745) | 2 | Tribe |

##### 1.2.2 Results

Simple updating of alignments for comparison to other tools.

The extension of the ITS *Senecio* alignment to include additional sequences of *Senecio* species, representatives of the different genera of the tribe Senecioneae and representatives of the different tribes within Asteroideae, resulted in an alignment comprising 703 sequences of 633 species that belong to 172 genera. Out of those, the genus *Senecio* (in its broad circumscription) is represented with 390 species.

In total there are 18,334 nucleotide sequences of 1,450 Senecioneae taxon ID's (including hybrids and subspecies) available in the local database. The taxonomy database holds information about 1,656 taxa. The updated alignment represents 1,154 species in total. Overall, PhylUp did not retain all taxa with sequences available on GenBank. Often this can be attributed to the fact that the loci I used, has no sequences for a particular taxon.

Across all alignments the sampling was increased after updating using PhylUp (Table 1, S2). According to Pelser et al. (2007) 152 genera belong to the tribe Senecioneae. Since then 10 new genera have been described – six with available sequences which are represented in the updated phylogeny (Cron, 2013; Dillon and Cruz, 2010; Hamzaoğlu et al., 2011; Liu and Yang, 2011;

Nordenstam, 2012; Nordenstam and Pelser, 2012; Nordenstam et al., 2009a; Pruski, 2012; Ren et al., 2017). One genus has since then been transferred to another genus within the tribe (Nordenstam et al., 2009b) and two genera are synonymized within the NCBI taxonomy: *Aetheolaena* – which needs to be transferred together with others to the genus *Senecio*. The representative sequence of *Cissampelopsis* (DC.) Miq. present in Pelser et al. (2007) is named 'Mikaniopsis sp. KAD343' in the NCBI taxonomy. One genus, which appears to be new in the updated phylogeny, *Cacalia* Kuntze, is actually a synonym of *Humbertacalia* C. Jeffrey. Overall, today belong 160 genera to the tribe. While in Pelser et al. (2007) 114 genera were represented by nuclear sequences, the updated phylogeny represents 148 genera, only for 12 genera are no sequences available yet.

The phylogenies produced in this study do not conflict with the support found in Pelser et al. (2007) and Pelser et al. (2010) (see 8.3). The support of the backbone of the nuclear phylogeny is similar to the ITS phylogeny by Pelser et al. (2007). In comparison to the concatenated ITS and ETS phylogeny by Pelser et al. (2010) which contains much less taxa, the different larger clades are less well supported in this study and support of clades within the genus *Senecio* is lower as in the respective publication by Kandziora et al. (2017). Both is likely a result of the much larger taxon and sequence sampling. The newly sampled genera all fall within clades that also have been supported by Cron (2013), Pelser et al. (2010), Pelser et al. (2007), and Ren et al. (2017).

Table S2: Total number of taxa present at different levels after alignment extension using PhylUp.

| **Loci** | **Genera** | **Species** | **Sequences** |
|---|---|---|---|
| ETS | 133 | 611 | 719 |
| ITS | 145 | 1157 | 1702 |
| *Trn*C-*ycf*6 | 24 | 323 | 430 |
| *Trn*LF | 137 | 648 | 803 |
| Nr | 149 | 1141 | 1345 |
| Cp | 138 | 750 | 832 |

Table S3: Number of sequences and taxa present in the different original single-marker alignments of Senecioneae and after extension using PhylUp.

| **Locus** | **ETS** | **ITS** | ***Trn*C-*ycf*6** | ***Trn*LF** |
|---|---|---|---|---|
| sequences in input data | 67 | 147 | 81 | 104 |
| species in input data | 67 | 146 | 75 | 95 |
| sequences added | 652 | 1555 | 349 | 699 |
| species added | 549 | 1054 | 300 | 590 |

#### Building of a customized sampled alignment

The single locus alignments for the Rosaceae phylogeny are well sampled and fulfill the requirements of the taxon sampling as set for the PhylUp run. The phylogeny based on the concatenated alignment is shown in Figure 1.1.

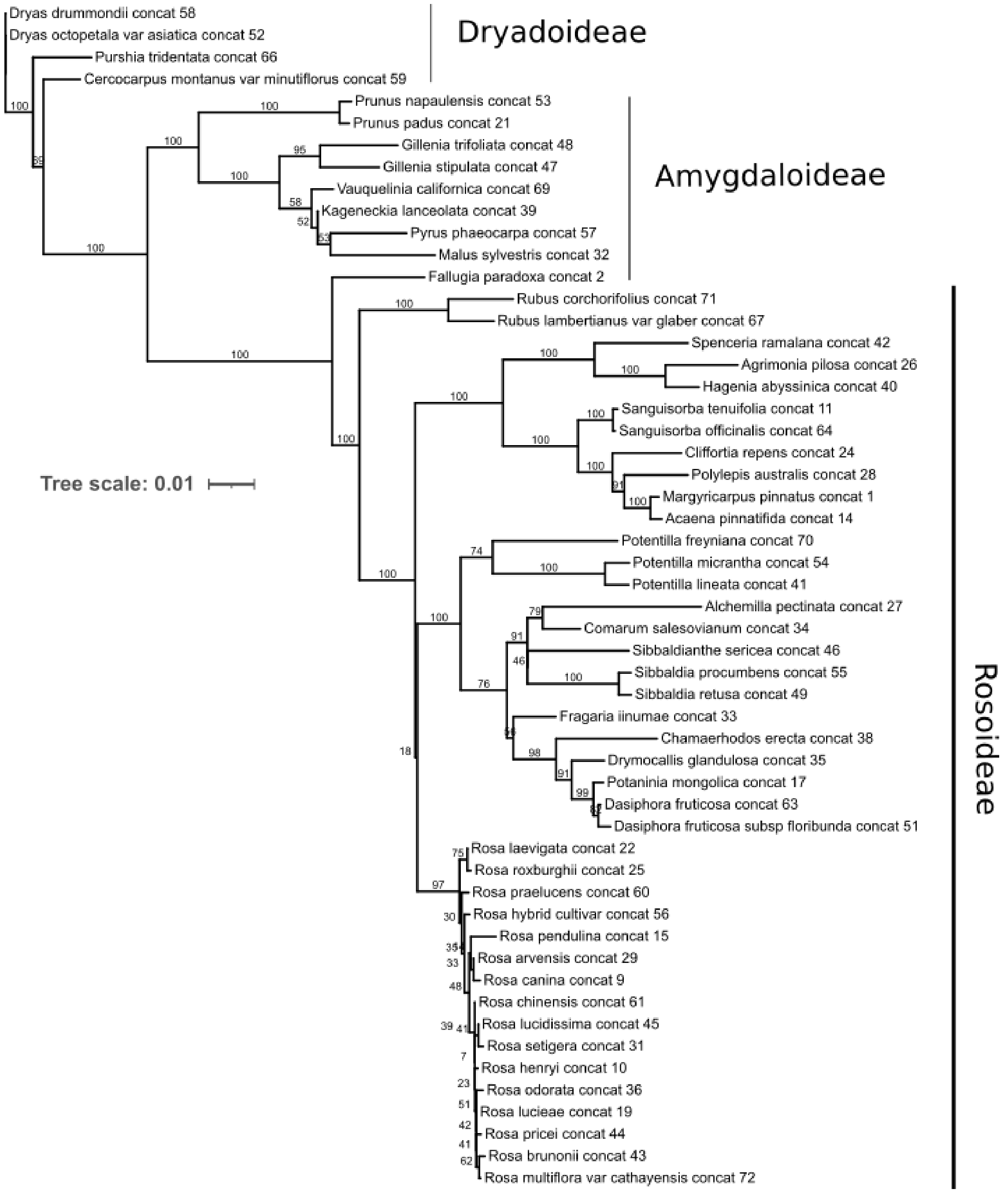


*Figure 1.1: Rosaceae plastid phylogeny calculated using RAXML-NG based on an alignment created using PhylUp.*

#### 1.3 Comparison of PhylUp to similar tools

##### 1.3.1 Similar programs and their settings

I compared the PhylUp results to two commonly used phylogeny reconstruction workflows: pyPHLAWD (Smith and Walker, 2019) and OneTwoTree (Drori et al., 2018). The OneTwoTree analysis was run from their website, which currently offers a beta-version (job was submitted on December 02, 2019). I chose the following settings, that need to be set or are different from the default: the ingroup was set to be Senecioneae, species descendants were set to exclude subspecies and to include hybrids, loci were chosen from the nuclear and chloroplast genomes and as phylogenetic inference tool, RAxML with 500 bootstrap iterations was selected.

The pyPHLAWD analysis relies on the master version from GitHub on December 3rd, 2019. First, I downloaded a new plant database using the phlawd db maker tool (https://github.com/blackrim/phlawd_db_maker; downloaded the master version on December 3rd, 2019 and the newest GenBank release (v. 233) for plants - which includes all plant and fungal sequences from before August 15, 2019). I then used that database to initiate a clustering run for the tribe Senecioneae.

##### 1.3.2 Comparison of Results

The alignments build using PhylUp returned more species than both, OneTwoTree and pyPHLAWD. Species sampling is maximized in PhylUp. Compared to PhylUp, the single marker alignments of the two tools are not trimmed at the ends which results in generally longer alignments than with PhylUp, but with more missing data (Table S4).The number of sequences and taxa present in the alignments produced by the different tools vary and the alignments of the same loci produced by the different tools have different species sampled and differ in the length of the alignments (see Tables S4 and S5).

With regard to the sampled loci, the Senecioneae analysis by OneTwoTree returns 15 different alignments and pyPHLAWD. The alignments with most sequences available in pyPHLAWD and OneTwoTree are the same loci as used with PhylUp. Both tools used for comparison have some alignments with similar description, e.g. *trn*K-*trn*K, trnL-*trn*F, *ndh*F. Assembling for example both *trn*L-*trn*F clusters by hand shows that one of the clusters overlaps with the second one. OneTwoTree finds more sequences for the *psb*A alignment, which is not well sampled in pyPHLAWD and not used in PhylUp.

Table S4: Number of species not represented across the different datasets. Total number of taxa represented by PhylUp is 1311, by OneTwoTree 1120 and by pyPHLAWD 1072.

| present/absent | PhylUp | OneTwoTree | PyPHLAWD |
|---|---|---|---|
| PhylUp | - | 290 | 328 |
| OneTwoTree | 99 | - | 218 |
| pyPHLAWD | 89 | 170 | - |

Table S5: Minimum, maximum and average sequence length of the different single-locus alignments using PhylUp, OneTwoTree and pyPHLAWD in comparison.

| Loci | PhylUp | | OneTwoTree | | pyPHLAWD | |
|---|---|---|---|---|---|---|
| | Mean | Min:max | Mean | Min:max | Mean | Min:max |
| ETS | 391.3408 | 249:497 | 416.56 | 265:556 | 414.24 | 330:494 |
| ITS | 630.0441 | 373:650 | 670.48 | 208:905 | 690.76 | 436:898 |
| *Trn*C-*ycf*6 | 724.6581 | 450:778 | 691.62 | 467:760 | 778.75 | 467:840 |
| *Trn*LF | 727.2055 | 394:818 | 762.18 | 321:951 | 834.4 | 406:944 |

##### 1.3.3 Discussion of PhylUp in comparison to similar tools

Comparing PhylUp to OneTwoTree and pyPHLAWD shows major differences. OneTwoTree and pyPHLAWD search all available loci in GenBank, while PhylUp only considers loci provided as input alignments - which also allows that carefully curated alignments by researchers can be used. Hence, the sampling of loci in PhylUp is lower. Nevertheless, the taxon sampling is maximized in PhylUp compared to the two other tools (Table S3). The alignments generated with the different tools have some taxa present that are missing in the other ones (Table S4). Some taxa are missing in PhylUp that are present in the alignments of the other two tools due to different reasons. First, due to our input requirement of a minimum sequence length, taxa that have only shorter sequences on GenBank were not added. For example, I used an alignment based on ITS1 and ITS2 as input, hence the sequence length requirement (65%) was too high to add sequences that only represent ITS1 or ITS2. This is likely also a reason why pyPHLAWD and OneTwoTree have several alignments for ITS and some chloroplast regions, which are added by OneTwoTree and pyPHLAWD as a shorter separate cluster. As such, the taxon sampling can be increased in PhylUp by permitting shorter sequences to be added (min_len, e.g. in the case of ITS: 0.35 adds ITS1/ITS2 sequences). Another solution would be to use ITS1 and ITS2 as separate input alignments. Second, some taxa are missing in PhylUp as those are only represented by loci not sampled by PhylUp.

Yet, taxa that are present in PhylUp are missing in OneTwoTree and pyPHLAWD. This is partially due to the usage of different GenBank databases (OneTwoTree: December 2018, pyPHLAWD: August 2019, PhylUp: October 2019). Many of the taxa that were not sampled by OneTwoTree and pyPHLAWD were added to GenBank before 2018, why those taxa are missing in alignments of pyPHLAWD or OneTwoTree is not obvious. Furthermore, in comparison to OneTwoTree and pyPHLAWD, the PhylUp alignments have much less missing data through the trimming of the alignment ends and deletion of short sequences (Table S5).

##### 1.3.4 References Supplement

Cron, GV (2013). “*Bertilia* — A new monotypic genus in the Senecioneae (Asteraceae) from South Africa”. In: South African Journal of Botany 88, pp. 10–16.

Dillon, MO and MZ Cruz (2010). “*Angeldiazia weigendii* (Asteraceae, Senecioneae), a new genus and species from northern Peru”. In: Arnaldoa 17.1, pp. 19–24.

Drori, M., Rice, A., Einhorn, M., Chay, O., Glick, L., & Mayrose, I. (2018). OneTwoTree: An online tool for phylogeny reconstruction. *Molecular ecology resources*, *18*(6), 1492-1499.

Hamzaoğlu, E, Ü Budak, and A Aksoy (2011). “A new genus, *Turanecio*, of the Asteraceae (tribe Senecioneae)”. In: Turkish Journal of Botany 35.5, pp. 479–508.

Kandziora, M, JW Kadereit, and B Gehrke (2016). “Frequent colonization and little in situ speciation in *Senecio* in the tropical alpine-like islands of eastern Africa”. In: American Journal of Botany 103.8, pp. 1483–1498.

– (2017). “Dual colonization of the Palaearctic from different regions in the Afrotropics by *Senecio*”. In: Journal of Biogeography 44.1, pp. 147–157.

Nordenstam, B (2012). “*Crassothonna* B. Nord., a new African genus of succulent Compositae-Senecioneae.” In: Compositae Newsletter 50, pp. 70–77.

Nordenstam, B and PB Pelser (2012). “*Caputia*, a new genus to accommodate four succulent South African Senecioneae (Compositae) species”. In: Compositae Newsletter 50, pp. 56–69.

Nordenstam, B, PB Pelser, and LE Watson (2009a). “*Lomanthus*, a new genus of the Compositae-Senecioneae from Ecuador, Peru, Bolivia and Argentina B Nordenstam, PB Pelser, LE Watson - Compositae Newsletter, 2009”. In: Compositae Newsletter 47, pp. 33–40.

– (2009b). “The South African aquatic genus *Cadiscus* (Compositae-Senecioneae) sunk in Senecio”. In: Compositae Newsletter 47, pp. 28–32.

Pelser, PB, AH Kennedy, EJ Tepe, JB Shidler, B Nordenstam, JW Kadereit, and LE Watson (2010). “Patterns and causes of incongruence between plastid and nuclear Senecioneae (Asteraceae) phylogenies”. In: American Journal of Botany 97.5, pp. 856–873.

Pelser, PB, B Nordenstam, JW Kadereit, and LE Watson (2007). “An ITS phylogeny of tribe Senecioneae (Asteraceae) and a new delimitation of *Senecio* L”. In: Taxon 56.4, pp. 1077–1104.

Smith, S. A., & Walker, J. F. (2019). PyPHLAWD: A python tool for phylogenetic dataset construction. *Methods in Ecology and Evolution*, *10*(1), 104-108.
